# FERMI: A novel method for sensitive detection of rare mutations in somatic tissue

**DOI:** 10.1101/208066

**Authors:** L. Alexander Liggett, Anchal Sharma, Subhajyoti De, James DeGregori

**Affiliations:** Department of Biochemistry and Molecular Genetics, University of Colorado School of Medicine, Aurora, CO 80045, USA.; Linda Crnic Institute for Down Syndrome, University of Colorado School of Medicine, Aurora, CO 80045, USA.; Rutgers Cancer Institute, New Brunswick, NJ 08901, USA.; Integrated Department of Immunology, University of Colorado School of Medicine, Aurora, CO 80045, USA.; Department of Pediatrics, University of Colorado School of Medicine, Aurora, CO 80045, USA.; Department of Medicine, Section of Hematology, University of Colorado School of Medicine, Aurora, CO 80045, USA.

## Abstract

With growing interest in monitoring mutational processes in normal tissues, tumor heterogeneity, and cancer evolution under therapy, the ability to accurately and economically detect ultra-rare mutations is becoming increasingly important. However, this capability has often been compromised by significant sequencing, PCR and DNA preparation error rates. Here, we describe FERMI (Fast Extremely Rare Mutation Identification) - a novel method designed to eliminate the majority of these sequencing and library preparation errors in order to significantly improve rare somatic mutation detection. This method leverages barcoded targeting probes to capture and sequence DNA of interest with single copy resolution. The variant calls from the barcoded sequencing data are then further filtered in a position-dependent fashion against an adaptive, context-aware null model in order to distinguish true variants. As a proof of principle, we employ FERMI to probe bone marrow biopsies from leukemia patients, and show that rare mutations and clonal evolution can be tracked throughout cancer treatment, including during historically intractable periods like minimum residual disease. Importantly, FERMI is able to accurately detect nascent clonal expansions within leukemias in a manner that may facilitate the early detection and characterization of cancer relapse.

## Background

The simultaneous growth in accuracy and reduction in cost of DNA sequencing has encouraged its use throughout many diverse areas of biology. Accompanying this explosion of applications for sequencing has been a natural demand for increasingly sensitive sequencing methods. While the detection of high frequency variants like germline SNPs is not particularly challenging by most sequencing technologies, sequencer and library preparation error rates are typically high enough to mask most rare or somatic variants. What is perhaps most challenging about library preparation is that the very isolation of DNA exposes it to oxidation that can change base identities (Shibutani, Takeshita, and Grollman 1991; Cheng et al. 1992), and high temperature exposure can thermally alter nucleotide identities (Lindahl and Karlstrom 1973; Lindahl and Nyberg 1974).

Because of these sequencing and library preparation limitations, quantitative PCR (qPCR) and multiparameter flow cytometry (MFC) have still remained common methods of rare variant detection (Terwijn et al. 2013). More recent technologies such as high-throughput digital droplet PCR (Hindson et al. 2011; Sykes et al. 1992; Vogelstein and Kinzler 1999), COLD-PCR (Milbury et al. 2012; J. Li et al. 2008), and BEAMing (Dressman et al. 2003) have shown promise for rare mutation detection, but are often limited to variant allele frequencies (VAFs) greater than 1 percent or are restricted to assaying only a few chosen mutations at a time.

A number of studies have sought improvements in sequencing technology accuracies by targeting and labeling small regions of genomic DNA such as sMIPs (Hiatt et al. 2013), paired strand collapsing (Kennedy et al. 2014) and other targeting methods (Thol et al. 2018; Mansukhani et al. 2018; Onecha et al. 2018). Some groups have also incorporated error correction methods to eliminate sequencing and PCR errors, like PELE-Seq (Preston et al. 2016), and error correcting enrichment processes (Schmitt et al. 2015). While these targeting and enrichment methods have certainly improved rare variant detection, they are still often limited to detecting variants that exist in at least 1% of a sample, or are limited to simultaneous detection of only a handful of variants.

Here, we describe a novel integrated genomic method that utilizes single molecule tagging and position specific background correction to push the limit of detection to variants existing in as little as 0.01% of a sample. Initial detection improvements come from the quantitative tracking ability of molecular barcodes that facilitate the elimination of the vast majority of sequencer and PCR amplification errors. Combined with paired-end sequence collapsing, consensus reads are produced that contain reduced numbers of false variants.

In a similar manner to previous methods (Chaudhuri et al. 2017; Young et al. 2016), we then experimentally derive a background of expected errors for each position within the consensus reads. As we know that sequence context impacts nucleotide stability (Benzer 1961; Gaffney and Keightley 2005; Lercher, Williams, and Hurst 2001; Nachman and Crowell 2000; Hwang and Green 2004), we use this background to correct our consensus reads based on probability density functions created for each assayed nucleotide position. We build on previous methods by then extensively characterizing the background error probabilities that generally occur in our sequencing library preparations. These characterizations were sufficiently comprehensive that in some cases we were able to eliminate all background variants, and only detect known mutations within a sample i.e. reaching 100% specificity.

One recent application of especially sensitive sequencing technologies is assaying and understanding clonal evolution within cancerous tissues (Greaves and Maley 2012). The rarity of somatic mutations, even within the clonally expanding pool of cells that exists within a tumor, has limited the observation of changes that can occur. Such an understanding would be valuable, as cancer therapies often leave behind a small number of cells that can frequently lead to relapse. In leukemias, the state during which these small numbers of cells remain after initial treatment is referred to as minimal residual disease (MRD). During this MRD stage, residual leukemia cells continue to evolve, and successful detection of relapsing leukemia at early stages may facilitate improved prognosis and treatment strategies (Ivey et al. 2016; Krönke et al. 2011).

As a proof of principle, we directly sample leukocyte genomic DNA and demonstrate the ability of FERMI to detect oncogenic changes during the MRD state, and monitor clonal changes with time. We also show that by concurrently sampling a diverse panel of oncogenic regions, we can detect the expansion of new oncogenic variants during MRD. Such observations could be critically important in predicting relapse in patients.

## Results

### Method Overview

We devised FERMI as a method to overcome current sequencing challenges facing rare mutation detection. FERMI is based on Illumina’s TrueSeq Custom Amplicon and AmpliSeq Myeloid protocols, which are designed for mutation detection across selected genomic regions. In FERMI, sequences found within human genomic DNA (gDNA) are captured by targeted oligomer probes, which are then sequenced and analyzed for the presence of any existing mutations. We adapted the AmpliSeq process to target a much smaller number of regions of the genome (32 vs 1500 regions) in order to achieve a greater sequencing depth per location with a reduced sequencing cost. We designed DNA probes to our 32 selected regions, each approximately 150bp in length, that span either AML-associated oncogenic mutations or Tier III (non-conserved, non-protein coding and non-repetitive sequence) regions of the human genome. The gDNA used for capture and sequencing was purified either from blood, cancer cell line or sperm cells, though most of our work focused on peripheral blood cells. The method should be adaptable to any species.

### Barcode-guided single molecule sequencing

Capture of gDNA, including any existing variants, begins by incubating double stranded gDNA together with oligomer probes designed to bind specified regions of the genome (Figure 1A). These probes span regions of approximately 150bp in length and contain two identifying indexes. The first index is a 16 bp sequence specific to each sample being processed, and the second is a 12 bp unique molecular identifier (UMI) of randomized DNA that should be unique to each captured strand of gDNA. Double stranded gDNA is melted apart to allow these targeting probes to bind the resulting single stranded DNA. Probe annealing is then achieved by slowly cooling the samples to allow for efficient targeting.

**Figure 1.**
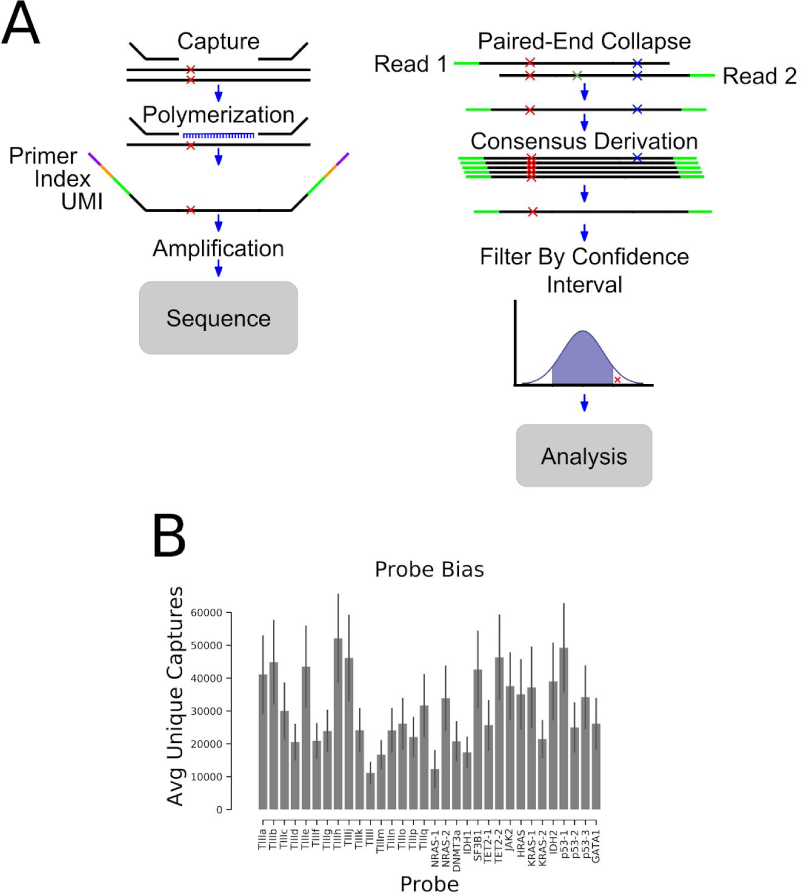
Method overview. A) Schematic representing the steps involved in identifying mutations with FERMI. B) The average number of unique captures varies by probe location. Error is standard deviation across 20 samples.

Following hybridization of the probes and gDNA, DNA polymerase is used to copy the template, and DNA ligase joins the strands together into a single contiguous amplicon. Using the sample indexes and the capture-specific UMIs to ensure each capture is tracked, amplicons are amplified by polymerase chain reaction (PCR) and pooled together for sequencing. Samples were sequenced using paired-end 150 bp sequencing, allocating approximately 30 million Illumina HiSeq or NovoSeq reads per sample. This coverage encompasses on average about 1,000,000 capture reactions per sample, resulting in about 30X sequencing coverage for each capture (given an average of 30,000 captures per probed region). Though capture efficiency was not uniform for the different probes, which show a 5-fold range in the numbers of successful unique captures, we show sufficient coverage at each probed location to capture mutations at least as rare as 0.01% (Figure 1B).

### Assessment of background error profile

Following sequencing, reads are distributed into sample-specific bins by their sample index. Within these sample-specific bins, paired-end reads are combined into single consensus reads by marking all mismatched base calls as an unknown identity. This approach yielded better results than elimination of pairs with some threshold of mismatches, as it retained substantially more sequencing information. These paired-end consensus reads are then sorted into capture-specific bins by their UMI sequences. These capture-specific bins are then collapsed into final consensus reads. In order to qualify for this final UMI-based consensus derivation, a UMI-specified capture is required to have at least 5 supporting sequencing reads, and the base at each position is only called if 75% of supporting reads agree with its identity. The final consensus reads are then compared against an experimentally determined background to distinguish true-positive variants from false positive signal, as described below.

Though UMI barcode collapsing of sequencing probes is an important technique by which sequencing sensitivity and accuracy can be increased (Hiatt et al. 2013), we find that UMI-collapsed data still retains a significant amount of false-positive variant signal. Using leukocyte gDNA purified from putatively healthy blood donors, we find approximately 5000 unique variants within our UMI collapsed consensus sequences in each individual. To estimate how much of this signal might be false-positive background, consensus sequences were computationally binned by the presence or absence of heterozygous SNPs found within our probed individuals. This sorting created bins of sequencing that should have originated from only a single allele. Theoretically, if rare variants were indicative of mutations that existed in-vivo, by their very nature of being rare, the mutations should exhibit an associative bias with only one of the two alleles. When we call variants within these two allele-specific bins however, we find that the variants associate quite uniformly across both alleles, suggesting that much of the variant signal found within our final consensus reads is erroneous (Figure 2A).

**Figure 2.**
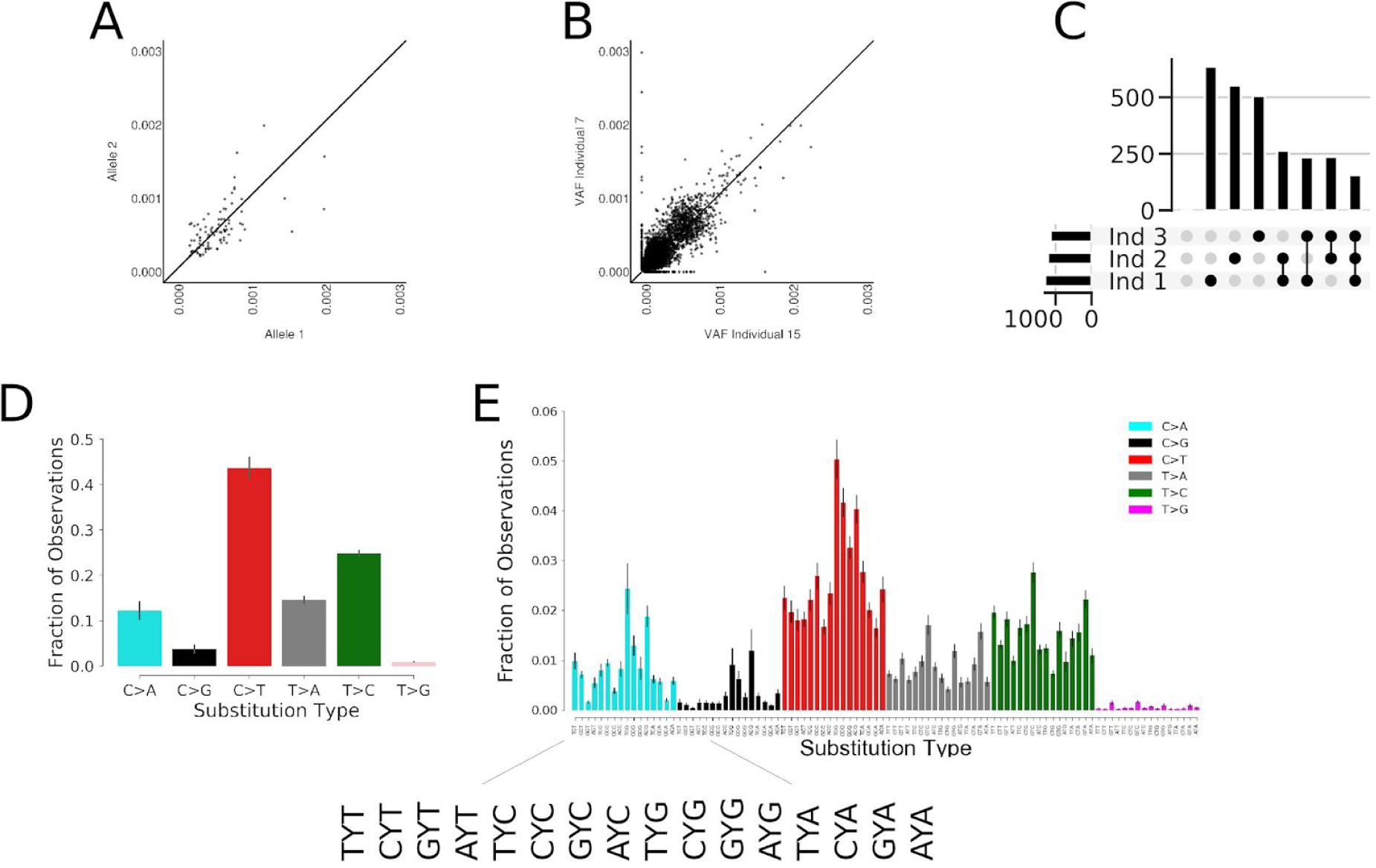
The background of false positive variants is similar across individuals. A) Using heterozygous SNPs to identify different alleles in human leukocyte gDNA, rare variants within consensus reads equally associate each of the alleles suggesting they are occurring randomly ex-vivo. B) Using FERMI to measure somatic mutation loads within two different samples shows that background signal is similar within leukocyte gDNA. C) The insertions/deletions found within three different individuals were identified. Indel counts are shown on the y-axis. Black dots represent sample groups, where the indel counts are those found within all samples indicated (either each sample alone, or commonly found across the indicated samples; for example, the next to the last vertical bar reflects indels found in both samples 3 and 2). Horizontal bars quantify the total number of indels found in a sample. D) Relative prevalence of observed substitutions within background signal found in leukocyte gDNA consensus reads. Complementary changes such as C>T and G>A are combined. Error is standard deviation across 20 individuals. E) Relative prevalence of observed substitutions classified by the neighboring upstream and downstream nucleotides (trinucleotide context). Error is standard deviation across 20 individuals.

Further suggestive of a significant false-positive presence within consensus reads, we show that when the rare variants found within the blood of any two individuals are compared, the same variants are found in each sample at nearly the same allele frequencies (Figure 2B). This similarity is not limited to inter-blood sample comparisons, as sperm gDNA shows similar patterns (Supplemental Figure 1). Finally, being similar to that in blood, the somatic mutation load we observe in sperm cells is well above previous estimates of less than 100 mutations per genome (Lynch 2016). Furthermore, when blood from healthy individuals was compared, mutations were no more similar in repeats from the same individual than between individuals (Supplemental Figure 4). Combined, these observations suggest that a false-positive background exists relatively uniformly across samples and sample types, and invites the possibility for a correction algorithm to distinguish real from false signal in order to significantly improve sequencing detection limits.

While over 90% of detected background variants were substitutions, occasionally insertions and deletions (indels) were observed. As many of these indels were observed in multiple captures for the same regions, we thought they might represent real mutations. However, as shown in Figure 2C, individual samples contain roughly 500 insertions or deletions, and about 250 of these are conserved across all samples. Furthermore, when a group of 20 individuals was pooled, only one insertion was not found at least twice within the pool. As these indels are often found in multiple captures, the repetitive occurrence between individual samples suggests that some mutagenic mechanism during sample processing is responsible for indel occurrence.

Within our final consensus reads, single-nucleotide substitutions account for the majority of falsely identified variants, and within this group of variants, there is significant identity bias. We find that C>T substitutions account for nearly 50% of the variants present in our final consensus reads, while other changes like C>G and T>G are far more rare (Figure 2D). Breaking down these substitutions into their trinucleotide contexts by including the bases located 5’ and 3’ of each change, we find that sequence context significantly impacts the probability of a false variant being identified (Figure 2E). Among the trinucleotide contexts, false variants within CpG sites are overrepresented within our final consensus reads.

Importantly, the patterns we identify in the trinucleotide context-independent and context-dependent substitutions mirror those identified in other studies of both normal tissues and cancers (Blokzijl et al. 2016; Martincorena et al. 2015; Alexandrov et al. 2013). The similarity of these patterns provides a cautionary note for mutation detection, as obedience to known patterns does not necessarily provide confidence in the accuracy of calls. It is possible that in-vivo mechanisms of mutation generation are similar to those experienced by template DNA ex-vivo, and therefore results in similar patterns within the background.

### Nucleotide context insufficiently explains background signal

In search of common patterns within our false positive background, we looked for surrounding sequence contexts that play a role in the prevalence of a false variant. While trinucleotide context does impact the probability that a substitution is found within our final consensus read pool, it often incompletely predicts the resulting variant allele frequency (VAF). We observe that many of the background substitutions found within our final consensus reads such as C>A within the contexts of CCA and ACA, exist within two relatively distinct VAF groups (Figure 3A). This indicates that within a given trinucleotide context, a substitution such as C>A will occur with either a high or a low frequency. Alternatively, some substitutions such as C>A within the CCG context largely occur with a low frequency, while other changes such as T>G in the context of CTA almost never exist at a high frequency. Both results suggest that trinucleotide context is not sufficient to predict background substitution rates at a given locus (Supplemental Table 1), consistent with recent reports that broader (epi)genomic contexts play key roles in replication errors, DNA damage, and repair (Coleman and De 2018). We do find, that regardless of the substitution identity or the trinucleotide context, a substitution defined only by trinucleotide context never exclusively occurs at high frequency.

**Figure 3.**
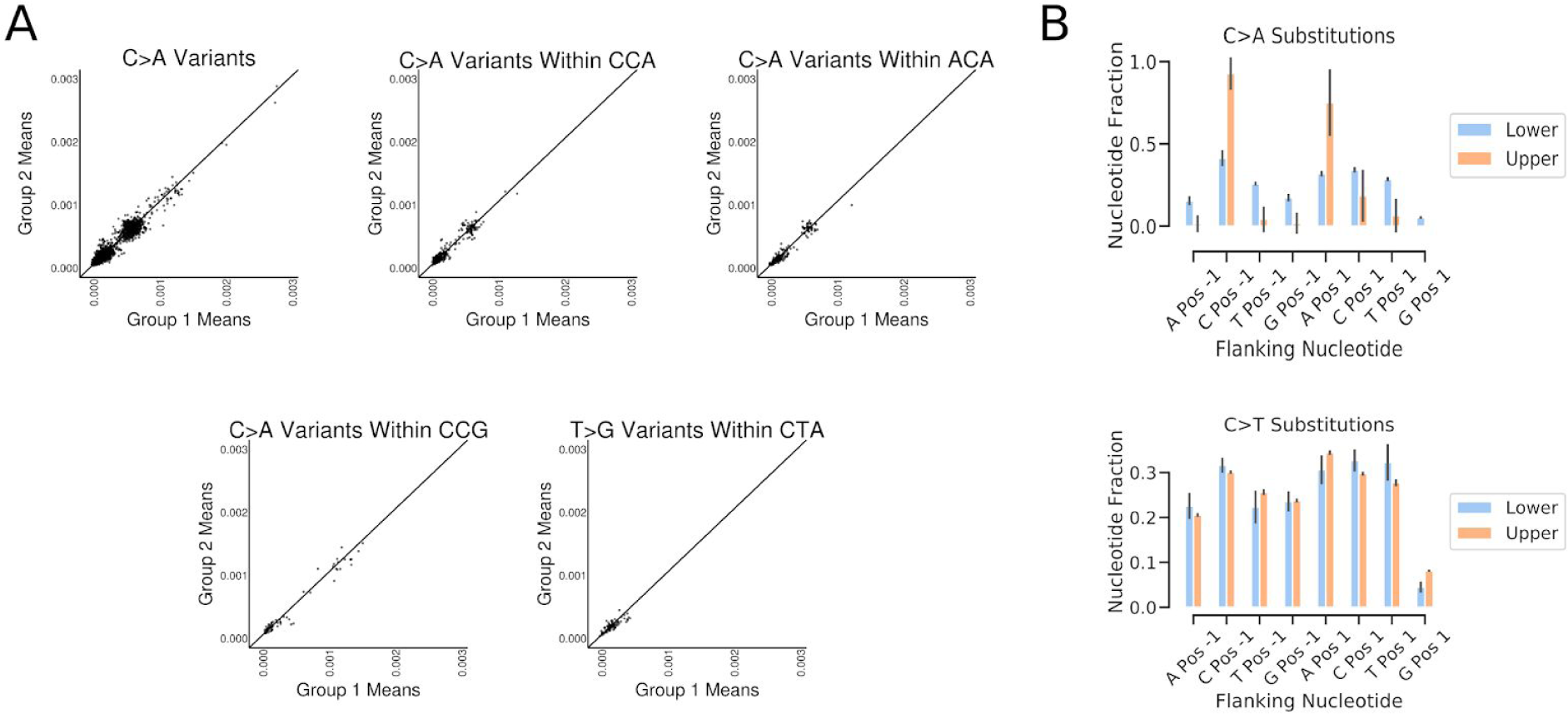
Trinucleotide context is insufficient to predict VAF. A) Mean VAF of background variants calculated from two different groups of 10 individuals probed with FERMI. Variants are either classified by substitution identity alone or within a particular trinucleotide context. B) Relative substitution rates for different substitutions classified by triplet context across all probed regions. Error is standard deviation across 20 individuals.

If the background variants are separated by their presence in either the upper or the lower VAF population, we find that for some changes such as C>A, both the 5’ and the 3’ nucleotides of the trinucleotide context significantly impact the VAF of the change (Figure 3B). This impact of the trinucleotide context is however not present for all changes, such as C>T substitutions, which show minimal bias of any of the possible trinucleotide contexts. We further searched for patterns within the 10bp upstream and downstream of a given change and find that only the triplet context showed any meaningful impact on mutation rate (Supplemental Figure 2).

### Algorithmic background subtraction eliminates most false positive signal

Although the trinucleotide context alone does not provide a sufficient amount of contextual information to determine the frequency with which a background variant is observed, nucleotide position strongly impacts the VAF of a substitution within the final consensus sequences. Throughout the probed regions, each nucleotide locus shows a unique background signal pattern that is relatively conserved across individuals (Figure 4A; similar conservation of background signal is observed across all other segments, data not shown). Some of the mutational patterns we observe within our background signal are similar to those found in The Cancer Genome Atlas (TCGA) (Supplemental Figure 3). Notably, enrichment for previously defined signatures are evident in background variants, representing artifacts of damage to isolated gDNA.

**Figure 4.**
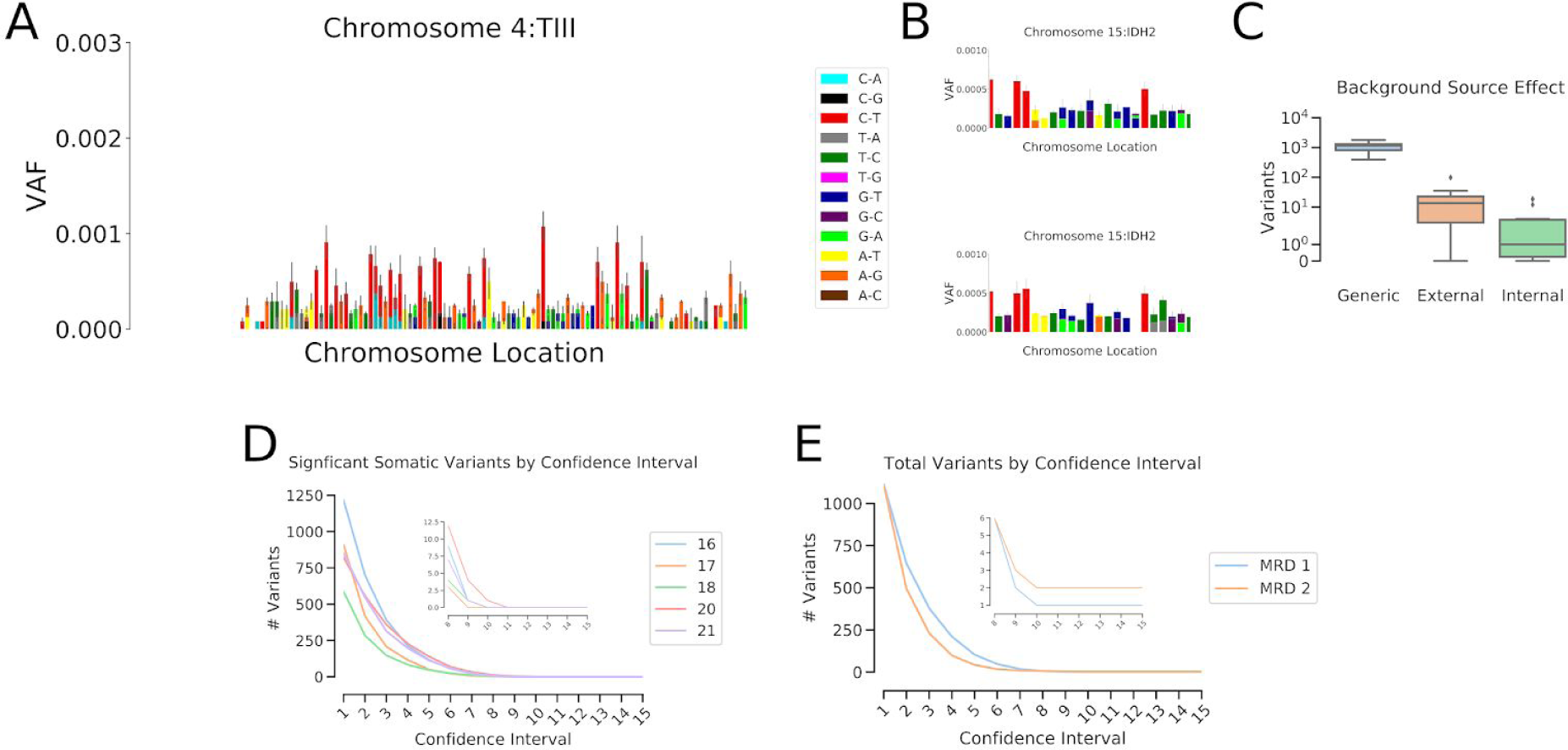
Background confidence intervals eliminate most variants. A) Observed substitution VAFs for a Tier III probe, illustrating the varying mutation presence at each nucleotide locus. Error is standard deviation across 20 individuals. B) Subsets of the IDH2 probe region from two different groups of 10 individuals illustrates the degree of similarity and differences that are commonly observed between samples. Error is standard deviation across the individuals. C) Total number of variants deemed significantly above background when only triplet context was used to generate expected background substitution rates (Generic), or when position-specific substitution rates generated from a different experiment (External) or the same experiment (Internal) are used. D) The total numbers of variants called as significantly above background for 5 individuals at confidence intervals from 0.9 - 0.999999999999999. E) The total numbers of variants called as significantly above background for two leukemic patients using sample 4 from MRD1 and sample 3 from MRD2 (See Figure 6), which are both points that had followed treatment, and at which leukemic burden was low.

Within our observed backgrounds, some nucleotide loci exhibit a strong bias towards a particular base change, showing only one type of substitution across all tested individuals. This effect is most commonly observed at nucleotides that exhibit a C>T substitution, where it is often the only observed change at that locus. Other nucleotide positions exhibit multiple different background substitutions, some changing to all three possible other bases. We noticed that while the background signal was surprisingly conserved across samples, that variability did exist (Figure 4B). It is often the case that a particular nucleotide locus will exhibit the same types of variants across different samples, but the allele frequencies vary.

Because each nucleotide locus tends to show a similar background across all tested individuals (Figure 2B), it was possible to derive a governing probability distribution for each observed substitution at every probed position. To create this distribution, a probability density function was created using a student’s T continuous random variable function. This probability density function was then used to calculate the high and low VAF endpoints of a confidence range by using a specified alpha fraction of the distribution.

We compared a number of different types of backgrounds to understand which best allowed us to eliminate false variants (Figure 4C). Initially a generic background was created by deriving a probability density function for each type of nucleotide change based on its neighbors (the three nucleotide patterns shown in Figure 2D), independent of genomic position. This was modestly effective at eliminating background variants from samples, as it reduced the total variant calls by about 80 percent (Figure 4C, “Generic”). By incorporating positional information and deriving a density function for each observed substitution at all genomic loci, about 99.9 percent of variant calls were eliminated, providing very clean sequencing data. Importantly, experimental variability seems to play a significant role in the accuracy of the background. While the same sample sequenced across multiple experiments generally shows a very similar background, experimental variability does appear (Supplemental Figure 4). While a background derived from samples taken from a different experiment (“External”) will result in about 10 variants being called as real in a given peripheral blood sample, a background created from samples run in the same experiment (“Internal”) will result in about 1-3 variants being called as real (Figure 4C). As expected variants are always retained, but total number of significant variants is minimized when using an internal background created from samples of the same experiment, internal backgrounds are used for all subsequent analyses.

To understand what alpha fraction of the probability density functions should be called as bona fide mutations, we used 10 healthy blood samples to derive a confidence interval range for each observed substitution across all probed nucleotide positions. These confidence intervals were then compiled into a comprehensive false-positive background against which experimental samples were then compared.

For 5 healthy blood samples, variants were called from their derived consensus sequences, and then compared against the comprehensive background. Within these 5 samples, variants were called as confidently above background if their VAFs were high enough to fall within the specific alpha of their governing probability density function. As expected, as the confidence interval alpha fraction was increased from 0.9 (One 9) to 0.999999999999999 (Fifteen 9’s) the number of variants called as confidently above background exponentially decreases (Figure 4D). This method eliminates nearly all background signal by confidence interval alpha fractions in the range of ten 9’s. Furthermore, at higher confidence intervals even germline variants are often eliminated for being too close to background, indicating excessive stringency. Peripheral samples taken from leukemic patients at different points during therapy were also tested, and show similar exponential decreases in confident variant calls, though the overall numbers of variants are higher than in healthy blood (Figure 4E).

### Assessing FERMI sensitivity and specificity

To help understand the specificity of FERMI in detecting only true mutations, Molm13 acute myeloid leukemia cells were expanded in vitro after passing them through bottlenecks of 1, 100, or 1 million cells. We show that many mutations can be observed within the cell cultures started from 100 cells, given that the 100-cell bottleneck should create clones at approximately 1% frequency each with occasional variants in our probed region (Figure 5A). Heterozygous mutations are expected at allele frequencies of 0.005 at 2N loci, and at lower VAFs if a variant falls in a region with greater ploidy. Indeed, most variants fall within this range. As expected, very few mutations are detected in gDNA isolated from the cell cultures started from 1 million cells, as most mutations will exist at rare allele frequencies (below our limit of detection). Similarly, we observed no mutations within the cultures initiated with single cells, consistent with the low odds that a mutation would occur in our probed region during the ∼14 cell divisions required to generate the 10,000 cell limit of detection. The absence of mutations detected in the cultures initiated with 1 cell each, but the presence of mutations within the cultures initiated with 100 cells, indicates that we have sufficiently limited false positive variants, but retained true mutations.

**Figure 5.**
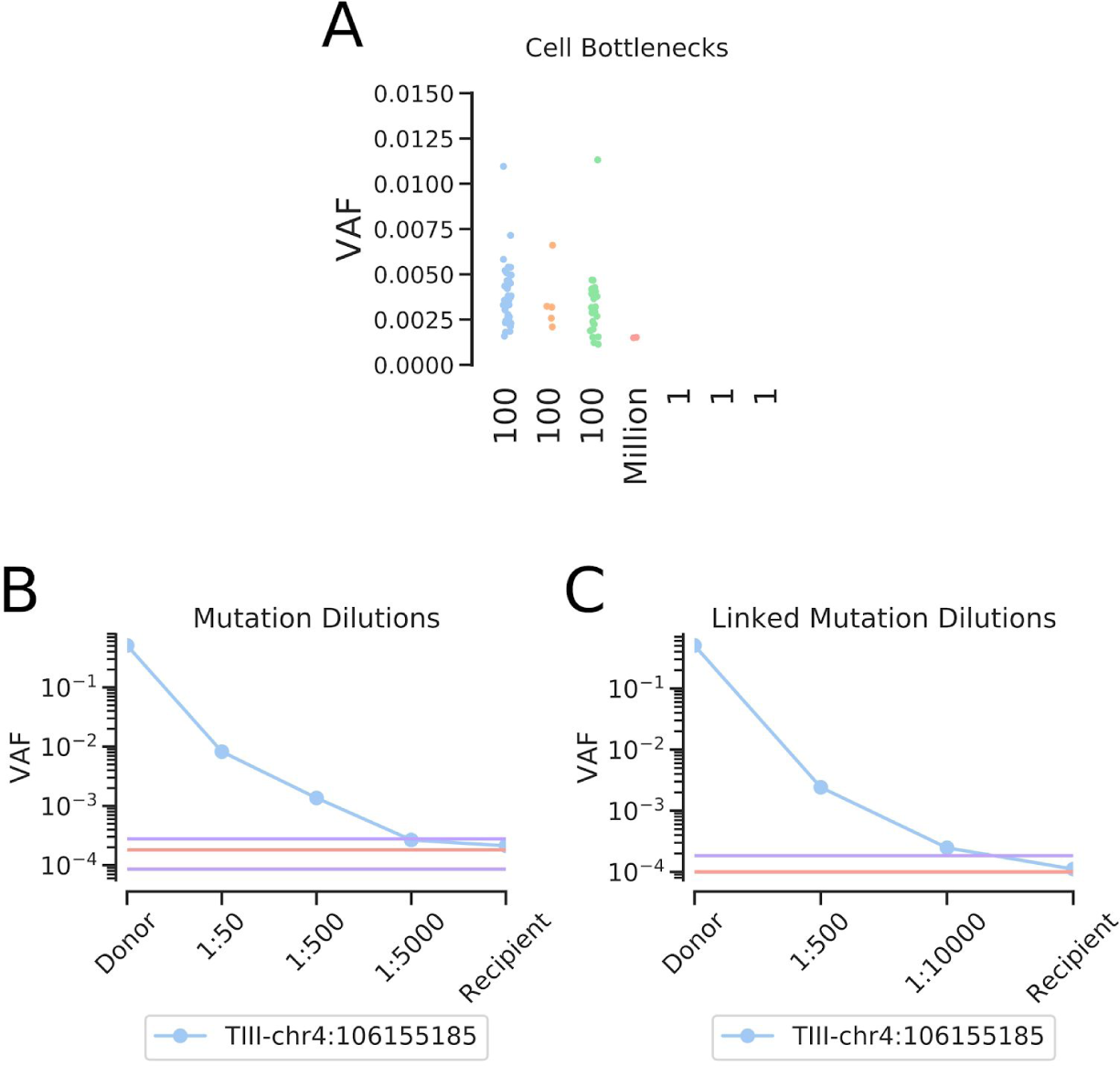
Assessing FERMI specificity and accuracy. A) MOLM-13 cells grown from 1, 100, or 1 million initial cells were expanded to a pool of 1 million cells, and then probed with FERMI. Samples illustrate how clonality impacts mutation detection by FERMI by altering the VAFs of somatic variants. B) Observed frequencies of a serially diluted blood gDNA sample with a heterozygous germline SNP show successful detection at allele frequencies from 1/2 to 1/10,000 (legend indicates the dilutions not the allele frequencies, where a 1/5,000 dilution of a heterozygous mutation should result in a 1/10,000 allele frequency). Background signal mean and standard deviation shown in red and purple respectively, calculated from 12 samples. C) Limit of detection improvements observed when multiple mutations in linkage disequilibrium are leveraged to eliminate erroneous reads. Background signal calculated from 12 samples.

To assess the sensitivity and limit of detection of FERMI, gDNA from human blood containing known heterozygous SNPs was serially diluted into blood gDNA lacking these SNPs. We find that the detection limits of tracking single dilutions to be variable as the level of background noise is position specific, but diluted germline variants were detected at frequencies at least as rare as 1:10,000 at the expected VAFs (Figure 5B). In other positions, where the background can be much higher, a dilution series would not be detected as low as 1:10,000.

One of the samples tested in the dilution series contained three heterozygous SNPs within the same probe region, on the same allele, allowing for an extra level of error correction. Within this sample, it was assumed that only those consensus reads with all three SNPs or those without any of the SNPs were correct, and all other reads were eliminated. This analysis significantly reduced the background error rates at the SNP positions, and allowed detection of diluted mutations at least as low as 1:10,000 (Figure 5C).

### Oncogenic driver detection in leukemias

To test the ability of FERMI to detect and follow mutations throughout leukemia treatments, patient biopsies were collected at disease inception, throughout treatment, and during relapse when possible. Using FERMI, we are able to detect these mutations when they are present and observe clonal evolution as it occurs. In some cases, we detect the principle oncogenic driver and watch it fluctuate in frequency in response to treatment without ever disappearing below background (Figure 6A). In another case, a JAK2 mutation is initially observed at high frequency, but treatment eliminates the clone. As relapse occurs, blast counts increase (Figure 6D,E,F), but the initial JAK2 clone does not increase in frequency, as the genetics of the leukemia has clearly changed with treatment (Figure 6B). In a third sample, we observe a patient relapse with a previously undetected JAK2 mutation (Figure 6C). While this time point was taken at relapse, we detect it at a frequency significantly below that of most sequencing method sensitivities, requiring only 5 ml of peripheral blood. The early detection of such a clone could allow treatment with a kinase inhibitor before overt disease relapse.

**Figure 6.**
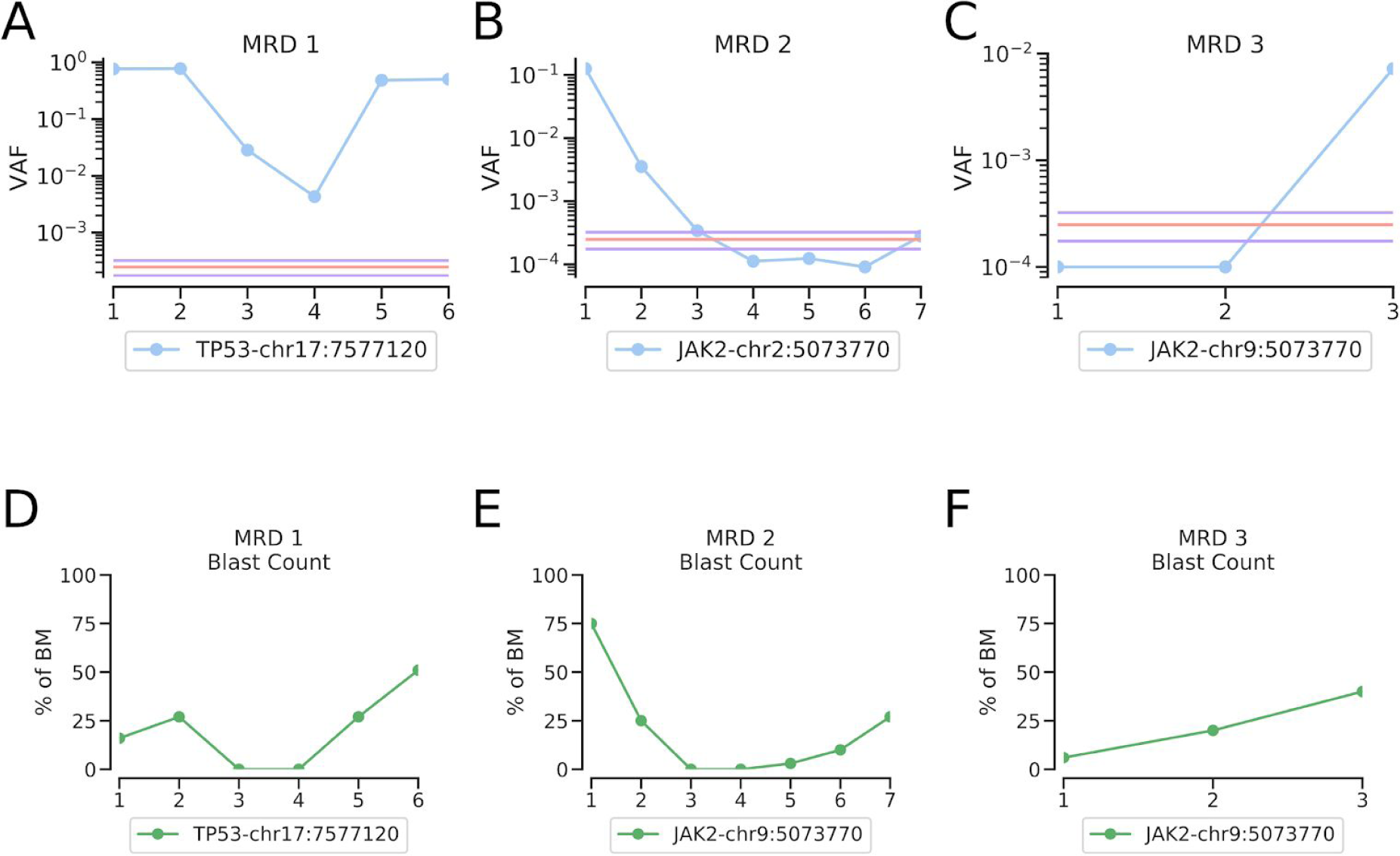
Oncogenic mutation detection during leukemia treatments. A) Oncogenic driver detection using FERMI on bone marrow biopsies taken at 6 different timepoints starting with clinical presentation of the patient and ending with relapse (x-axis is in chronological order of leukemia samplings). Background signal mean and standard deviation shown in red and purple respectively, and is derived from 20 samples. B) Oncogenic driver detection throughout leukemia treatment in a case where relapse was not driven by a mutation within our panel. Background derived from 20 samples. 27. C) Example of a leukemic relapse in which the detected driver was not unknown prior to blood sampling. Background derived from 8 samples. D,E,F) Corresponding blast counts as percentage of bone marrow biopsies.

### Discussion ∼750 words

In this study, we designed a sensitive sequencing method that enables the accurate detection of rare variants and clonal evolution within primary samples. In leveraging the quantitative power of capture-unique UMI barcodes, we achieve single-allele sequencing resolution from gDNA, and by then combining these sequencing results with a comprehensive analysis of expected background signal, we achieve exceptional sequencing fidelity.

While capture-specific barcoding has been effectively used in the past, an inability to achieve sufficient capture numbers and high background have often held the theoretical limit of detection to variants existing at a frequency of >1/1,000. By probing roughly 30,000 different unique captures for each region of interest per sample, we pushed our theoretical limit of detection to at least 1/10,000 (at least for positions where background signal is sufficiently low).

Paired-end collapsing has been successfully used to reduce the number of sequencing errors within sequencing data. Unfortunately, errors also occur during library preparation, and while the molecular barcodes assist with the elimination of these library preparation errors, mistakes made before or during the first round of PCR will typically appear indistinguishable from a heterozygous variant, such that neither paired-end collapsing nor molecular barcode collapsing will be capable of eliminating them. This understanding prompted the development of an expected false positive background that could be used to filter out common mistakes that occur during library preparation.

Our experimentally derived backgrounds proved vitally important in determining whether or not a variant found in a sample was sufficiently elevated in allele frequency that it could be classified as truly existing in the in-vivo gDNA from which it originated. Similar to background subtraction employed by other groups (Chaudhuri et al. 2017), our generalized background allowed us to not only detect expected mutations, but also discover new mutations within samples.

It is interesting to note that within our correction background, we observe similar mutation patterns to those observed for other studies, and even similar mutational signatures (Behjati et al. 2014; Alexandrov et al. 2013). This may indicate a surprising degree of similarity between the intrinsic mutagenic processes in-vivo and error-causing processes involved in sample preparation. It is possible that this similarity is the result of the conserved behavior of error-inducing machinery like DNA polymerase and DNA ligase both in-vivo and in-vitro, or even similar mutagenic exposures such as oxidative damage. These observed similarities suggest caution against using mutation signatures as validation of the accuracy of sequencing data when attempting to identify rare variants.

The early detection of clonal evolution within cancer samples has held the promise of more comprehensive diagnoses and improved treatment strategies for patients. While deep sequencing has been applied to patient leukemias in the past, mutation discovery accuracies have typically limited these approaches to more of a validation role (Thol et al. 2018). We obtained a number of leukemic patient biopsies, taken at initial clinical presentation, and throughout treatment, and used FERMI to search for somatic mutations. We find that while the detection limit of FERMI is quite low, the greatest improvements are made through its accurate mutation detection ability. Because we eliminate nearly all background variants, we can accurately detect unexpected relapse mutations and drivers of clonal expansions.

In requiring only around 5ml of blood, FERMI could be easily used in a clinical setting to quickly, cheaply and easily identify important driver mutations and clonal evolution within patient’s cancers. If relapse mutations were caught by FERMI when they are still rare, targeted therapies could be used to prevent them from clonally expanding to fixation and driving leukemic relapse.

## Acknowledgments

We would like to thank Ruth Hershberg of Technion University and Jay Hesselberth and Robert Sclafani of the University of Colorado School of Medicine for useful suggestions and for review of the manuscript. We would like to thank Craig Jordan and Amanda Winters of the University of Colorado School of Medicine for the primary leukemia samples. We would like to thank the Illumina Concierge team who assisted with the initial probe design. We would also like to thank Joe Hiatt, Beth Martin, and Jay Shendure of the Shendure lab, who assisted in the initial design of the probing technique. These studies were supported by grants from the National Cancer Institute (R01CA180175 to J.D.), NIH/NCATS Colorado CTSI Grant Number UL1TR001082CU (seed grant to J.D.), a SCOR grant from the Leukemia and Lymphoma Society (to Craig Jordan), F31CA196231 (to L.A.L.), the Linda Crnic Institute for Down Syndrome (to J.D. and L.A.L.), and P30-CA072720 (to A.S. and S.D.). The research utilized services of the Cancer Center Genomics Shared Resource, which is supported in part by NIH grant P30-CA46934. L.A.L. and J.D. developed the concept of this project, planned the experiments, analyzed results, and wrote the manuscript. L.A.L. processed and prepared samples from blood biopsy to sequencing, and wrote the software used for analysis. A.S. and S.D. analyzed results, and contributed to writing of the manuscript.

## Supplementary Materials

### Materials and Methods

#### Amplicon Design

Amplicon probes for targeted annealing regions were created using the Illumina Custom Amplicon DesignStudio (https://designstudio.illumina.com/). UMIs were then added to the designed probe regions and generated by IDT using machine mixing for the randomized DNA. Probes were PAGE purified by IDT. All probes are listed below along with binding locations and expected lengths of captured sequence.

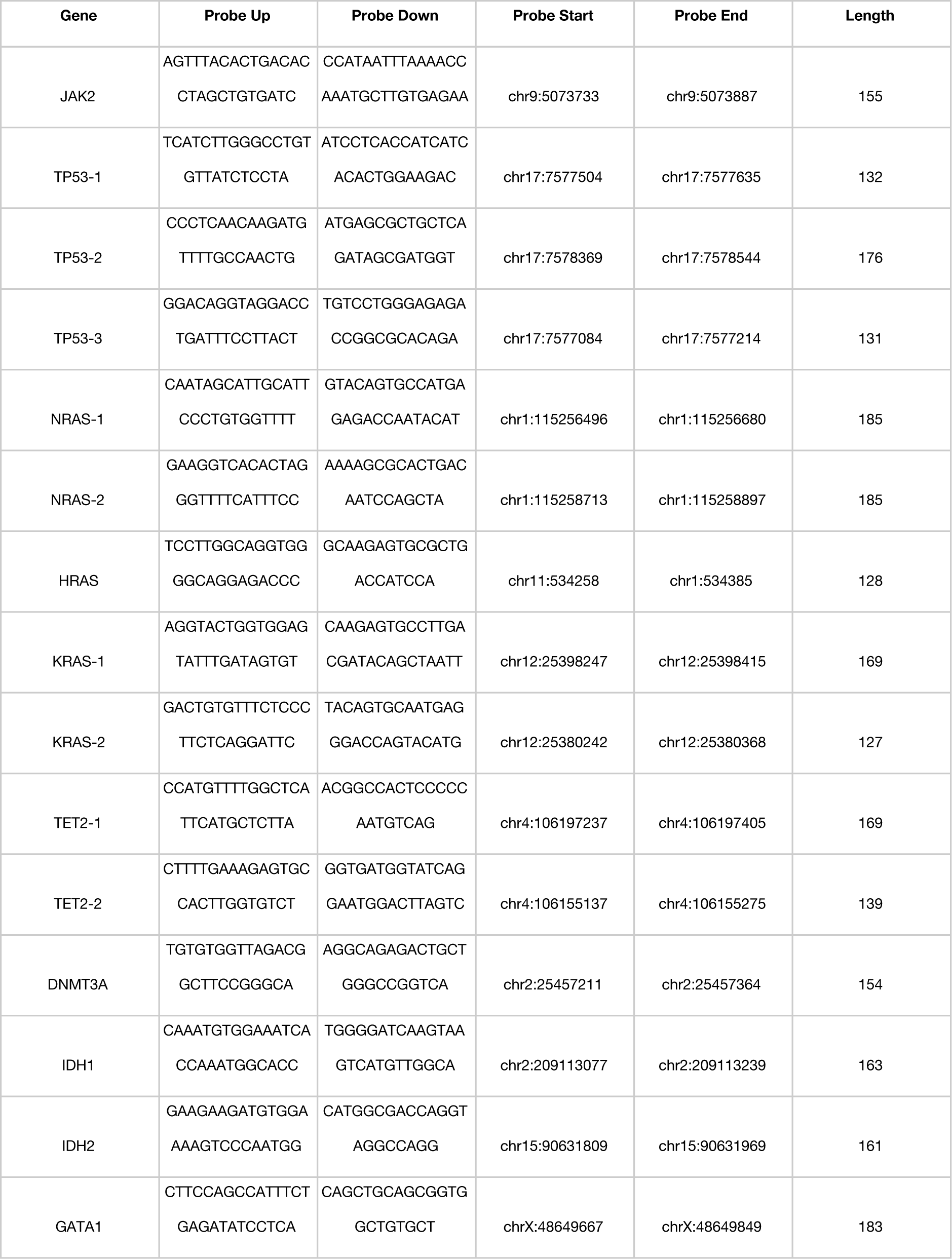
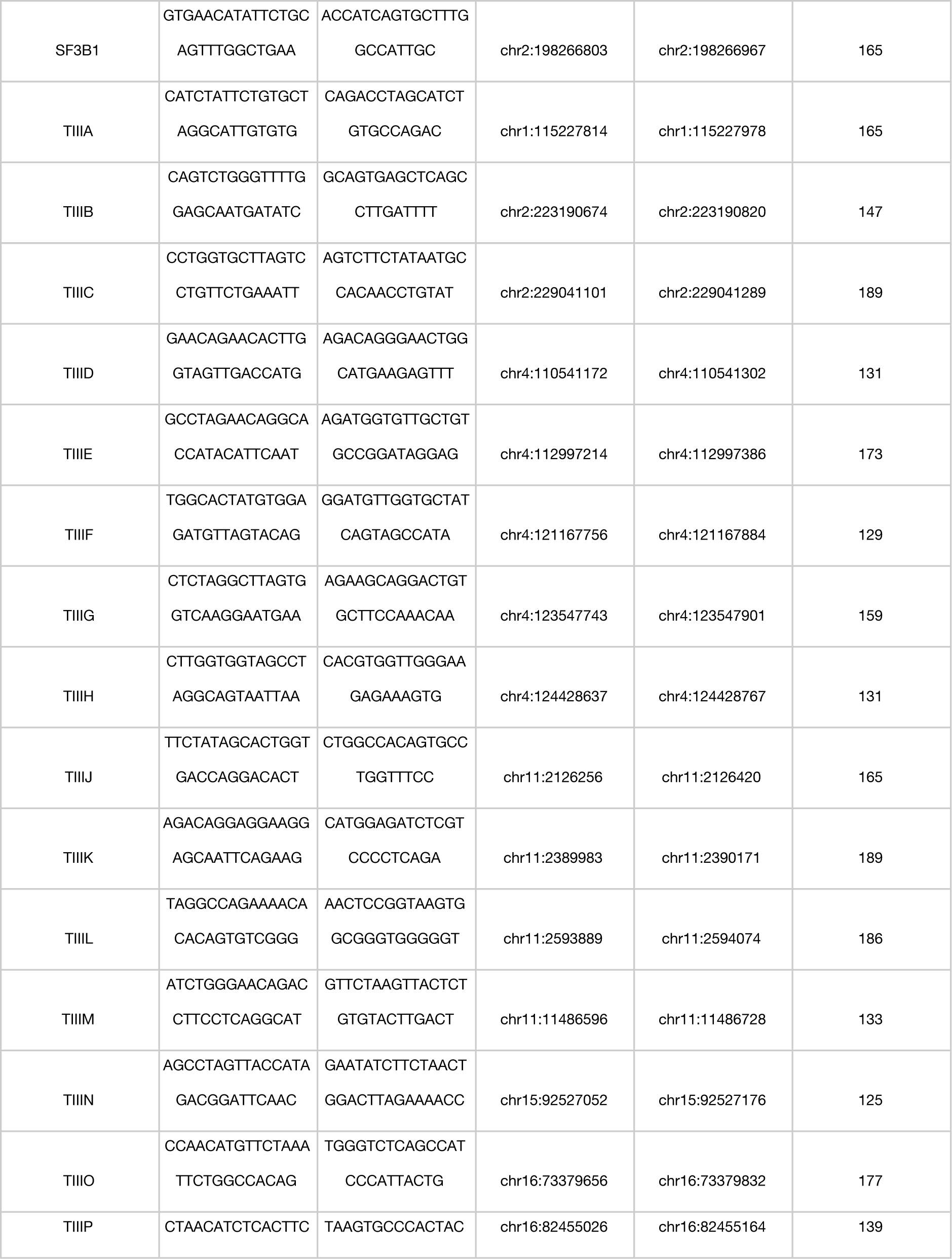
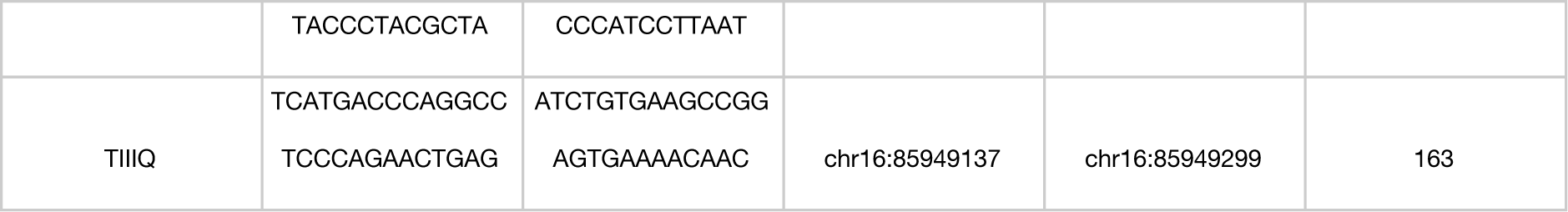

#### Genomic DNA Isolation

Human blood samples were purchased from the Bonfils Blood Center Headquarters of Denver Colorado. Our use of these deidentified samples was determined to be “Not Human Subjects” by our Institutional Review Board. Biopsies were collected as unfractionated whole blood from apparently healthy donors, though samples were not tested for infection. Samples were approximately 10 mL in volume, and collected in BD Vacutainer spray-coated EDTA tubes. Following collection, samples were stored at 4°C until processing, which occurred within 5 hours of donation. To remove plasma from the blood, samples were put in 50 mL conical tubes (Corning #430828) and centrifuged for 10 minutes at 515 rcf. Following centrifugation, plasma was aspirated and 200 mL of 4°C hemolytic buffer (8.3g NH_4_Cl, 1.0g NaHCO_3_, 0.04 Na_2_ in 1L ddH_2_O) was added to the samples and incubated at 4°C for 10 minutes. Hemolyzed cells were centrifuged at 515 rcf for 10 minutes, supernatant was aspirated, and pellet was washed with 200 mL of 4°C PBS. Washed cells were centrifuged for at 515rcf for 10 minutes, from which gDNA was extracted using a DNeasy Blood & Tissue Kit (Qiagen REF 69504).

#### Amplicon Capture

For amplicon capture from gDNA, we modified the Illumina protocol called “Preparing Libraries for Sequencing on the MiSeq” (Illumina Part #15039740 Revision D). DNA was quantified with a NanoDrop 2000c (ThermoFisher Catalog #ND-2000C). 500ng of input DNA in 15μl was used for each reaction instead of the recommended quantities. In place of 5μl of Illumina ‘CAT’ amplicons, 5μl of 4500ng/μl of our amplicons were used. During the hybridization reaction, after gDNA and amplicon reaction mixture was prepared, sealed, and centrifuged as instructed, gDNA was melted for 10 minutes at 95°C in a heat block (SciGene Hybex Microsample Incubator Catalog #1057-30-O). Heat block temperature was then set to 60°C, allowed to passively cool from 95°C and incubated for 24hr. Following incubation, the heat block was set to 40°C and allowed to passively cool for 1hr. The extension-ligation reaction was prepared using 90 μl of ELM4 master mix per sample and incubated at 37°C for 24hr. PCR amplification was performed at recommended temperatures and times for 29 cycles. Successful amplification was confirmed immediately following PCR amplification using a Bioanalyzer (Agilent Genomics 2200 Tapestation Catalog #G2964-90002, High Sensitivity D1000 ScreenTape Catalog #5067-5584, High Sensitivity D1000 Reagents Catalog #5067-5585). PCR cleanup was then performed as described in Illumina’s protocol using 45 μl of AMPure XP beads. Libraries were then normalized for sequencing using the Illumina KapaBiosystems qPCR kit (KapaBiosystems Reference # 07960336001).

#### Sequencing

Prepared libraries were pooled at a concentration of 5 nM. Libraries were sequenced on the Illumina HiSeq 4000 at a density of 12 samples per lane with 5% PhiX DNA included, or on the Illumina NovaSeq 6000, allocating approximately 30 million reads per sample.

#### Bioinformatics

The analysis pipeline used to process sequencing results can be found under FERMI here: http://software.laliggett.com/ or here: https://github.com/liggettla/FERMI. For a detailed understanding of each function provided by the analysis pipeline, please refer directly to the software. The overall goal of the software built for this project is to analyze amplicon captured DNA that is tagged with equal length UMIs on the 5’ and 3’ ends of captures, and has been paired-end sequenced using dual indexes. Input fastq files are either automatically or manually combined with their paired-end sequencing partners into a single fastq file. Paired reads are combined by eliminating any base that does not match between Read1 and Read2, and concatenating this consensus read with the 5’ and 3’ UMIs. A barcode is then created for each consensus read from the 5’ and 3’ UMIs and the first five bases at the 5’ end of the consensus. All consensus sequences are then binned together by their unique barcodes. The threshold for barcode mismatch can be specified when running the software, and for all data shown in this manuscript one mismatched base was allowed for a sequence to still count as the same barcode. Bins are then collapsed into a single consensus read by first removing the 5’ and 3’ UMIs. Following UMI removal, consensus sequences are derived by incorporating the most commonly observed nucleotide at each position, so long as the same nucleotide is observed in at least a specified percent of supporting reads (75% of reads was used for results in this manuscript) and there are least some minimum number of reads supporting a capture (5 supporting reads was used for results in this manuscript). Any nucleotide that does not meet the minimum threshold for read support is not added to the consensus read, and alignment is attempted with an unknown base at that position. From this set of consensus reads, experimental quality measurements are made, such as total captures, total sequencing reads, average capture coverage, and estimated error rates. Typically we required 5 total captures of a variant to be observed for the variant to be counted as real.

Derived consensus reads are then aligned to the specified reference genome using Burrows-Wheeler (H. Li and Durbin 2009), and indexed using SAMtools (H. Li et al. 2009). For this manuscript consensus reads were aligned to the human reference genome hg19 (Lander et al. 2001; Fujita et al. 2010) (though the software should be compatible with other reference genomes). Sequencing alignments are then used to call variants using the Bayesian haplotype-based variant detector, FreeBayes (Garrison and Marth 2012). Identified variants are then decomposed and block decomposed using the variant toolset vt (Tan, Abecasis, and Kang 2015). Variants are then filtered to eliminate any that have been identified outside of probed genomic regions. If necessary, variants can also be eliminated if below certain coverage or observation thresholds such that variants must be independently observed multiple times in different captures to be included.

The final variants called from the consensus sequences were then compared to experimentally derived confidence intervals for each probed position. These confidence intervals were created by using FERMI to sequence control peripheral blood samples from the same experiment as test samples. Following the logic described in Results, it was assumed that low frequency variants that are detected across multiple individuals (including in blood and sperm, where few variants are expected) were not real signal but rather false positive background. All of the variants from these control samples were thus used to construct a standard background. This background was calculated for each position at which a variant was observed within the standard control samples, and was uniquely calculated for each type of change. Often, in the construction of the background, the highest frequency alleles were eliminated in an effort to minimize the effect of true mutations on the background. A student’s T continuous random variable function was used to create a probability density function that describes the background distribution for each substitution type at every probed locus (Oliphant 2007). By specifying a particular alpha fraction of the distribution, high and low VAF endpoints were derived that were then used to determine if an experimental signal was significantly above background.

#### Elimination of false positive sequencing and library creation artifacts

A number of steps have been included within sample preparation and bioinformatics analysis specifically to reduce false background signal. Using the dilution series shown in Figures 1C-D, we can show sufficient sensitivity to identify signal diluted to levels as rare as 10^−4^. While these dilutions show significantly improved sensitivity over many current sequencing methods, background error could still exist. The two largest sources of erroneous mutation when sequencing DNA will typically be from PCR amplification mutations (caused both by polymerase errors and exogenous insults like oxidative damage), and sequencing errors.

These are the steps taken to eliminate errors before final background derivation:

- Elimination of first round PCR amplification errors
- Elimination of subsequent PCR amplification errors
- Elimination of sequencing errors

#### Elimination of first round PCR amplification errors in consensus reads

The first round of PCR amplification performed during library preparation causes mutations that are challenging to distinguish from those that occurred endogenously. Since there is little difference between those mutations that occur during the first round of PCR amplification and those that occurred endogenously, we rely on probability to eliminate these errors. Since we are performing sequencing of individually captured alleles, we can ask whether requiring that a mutation be observed in multiple captured alleles before it is called as a true positive signal alters the frequency of variants identified. We expect about 400 first round PCR amplification errors, and the probability that the identical mutation will occur in multiple cells becomes exponentially unlikely. By requiring a mutation be observed in just five captures before it is called as real signal, theoretically, none of the first round PCR amplification errors should make it into the final consensus reads.

#### Elimination of subsequent PCR amplification errors

Elimination of PCR amplification errors after the first round of PCR is done using UMI collapsing (Figure 1A). Each time a strand is amplified, the UMI will keep track of its identity. Any mutations that occur after the first round of PCR will be found on average in 25% of the reads (or fewer for subsequent rounds). This allows us to collapse each unique capture and eliminate any rarely observed variants (<75%) associated with a given UMI. Utilizing the UMI in this way allows us to essentially eliminate any PCR amplification errors that occurred after the first round of PCR. The method should also eliminate most errors resulting from DNA oxidation in vitro.

#### Elimination of sequencing errors

Sequencing errors are eliminated in two ways. This first method is by using paired-end sequencing to read each strand of a DNA fragment (Figure 1A). The sequence of these reads (Read1 and Read2) should match if no sequencing errors have been made. For an error to escape elimination it would need to occur at the same position (changing to the same new base) within both Read1 and Read2. Therefore, when the base call differs at a position on Reads 1 and 2, these changes are eliminated from the final sequence. This collapsing should eliminate most sequencing errors, although sequencing errors of the same identity occurring at the same position will escape. These errors should be removed when collapsing into single capture bins (Figure 1A). As with the logic when eliminating subsequent PCR amplification errors, most sequences associated with each UMI pair should be identical. Therefore, sequencing errors passing through Read1 and Read2 will be very unlikely to match other sequenced strands from the same capture event, and are eliminated during consensus sequence derivation.

## Supplemental Figures

**Supplemental Figure 1.**
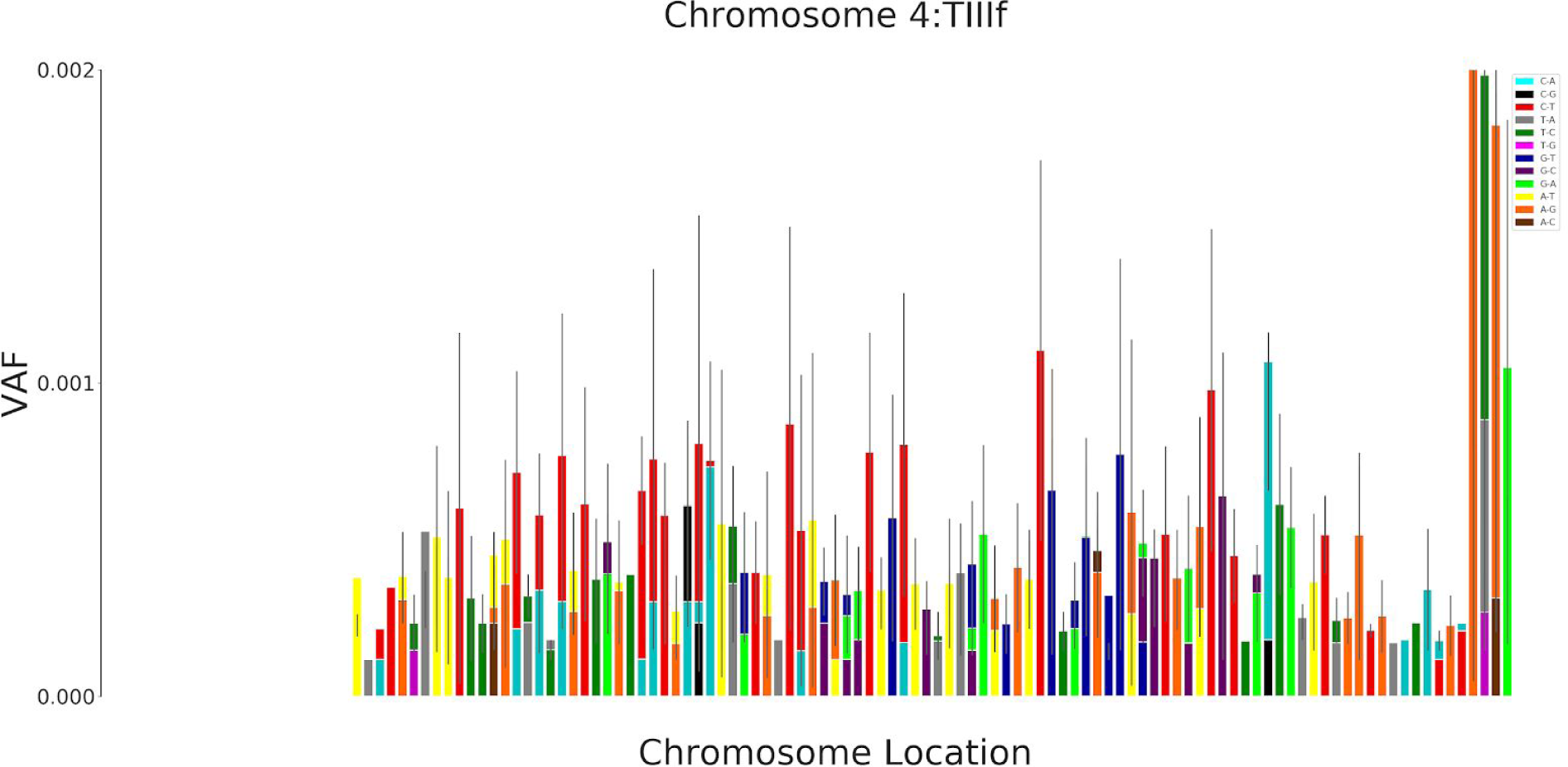
Sperm gDNA shows similar background to blood samples. Use of FERMI to sequence 12 samples of sperm gDNA shows a significant signal within uncorrected consensus reads. Work from other groups robustly shows that any given sperm cell should contain about 100 mutations, confirming the presence of a widespread background signal within our uncorrected consensus reads (Lynch 2016).

**Supplemental Figure 2.**
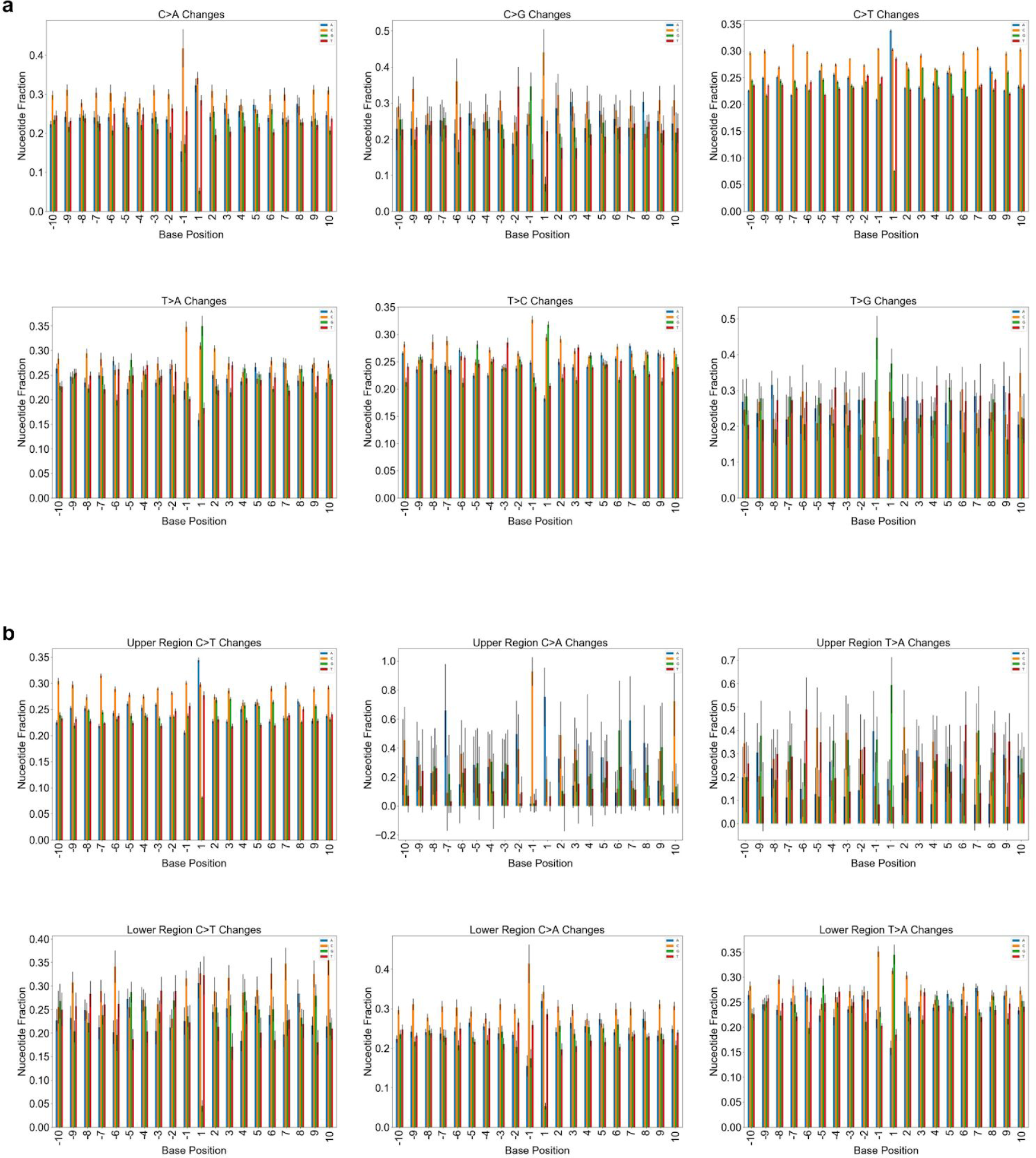
Trinucleotide sequence is more impactful on background substitution type than more distant surrounding bases. **a**, Shown for each of the 6 possible substitution types are the relative frequencies at which each of the various nucleotides are found at a given position surrounding the variant nucleotide (plotted as means with standard deviation between individuals as error; n = 22). Within each of the subplots, the variant is not shown and exists at position 0. It appears that the greatest skewing of nucleotide representation occurs at positions −1 and 1, suggesting that they have the greatest impact on how a base will change when it suffers a substitution in vitro. Note that for C changes, underrepresentation of G at position 1 is expected based on low representation of CpGs in the captured regions. **b**, As seen in Fig 2a-b, substitutions tend to exist within an upper or lower region of allele frequencies. To understand if flanking nucleotide sequence plays a role in this, the populations were analyzed separately for each of 6 base changes at Cs and Ts. Suggesting that the actual mutation plays a role in the resulting VAF, most substitutions exist disproportionately in either the upper or lower population rather than being equally distributed between the two. For some comparisons, this resulted in larger error within one of the populations, rendering some comparisons not feasible. For C>T changes, the flanking base sequence was largely conserved between the two populations. Other substitutions show differences in flanking sequence when they exist at higher or lower VAFs, as observed for T>A at positions −1 and 1 and C>A at position 1.

**Supplemental Figure 3.**
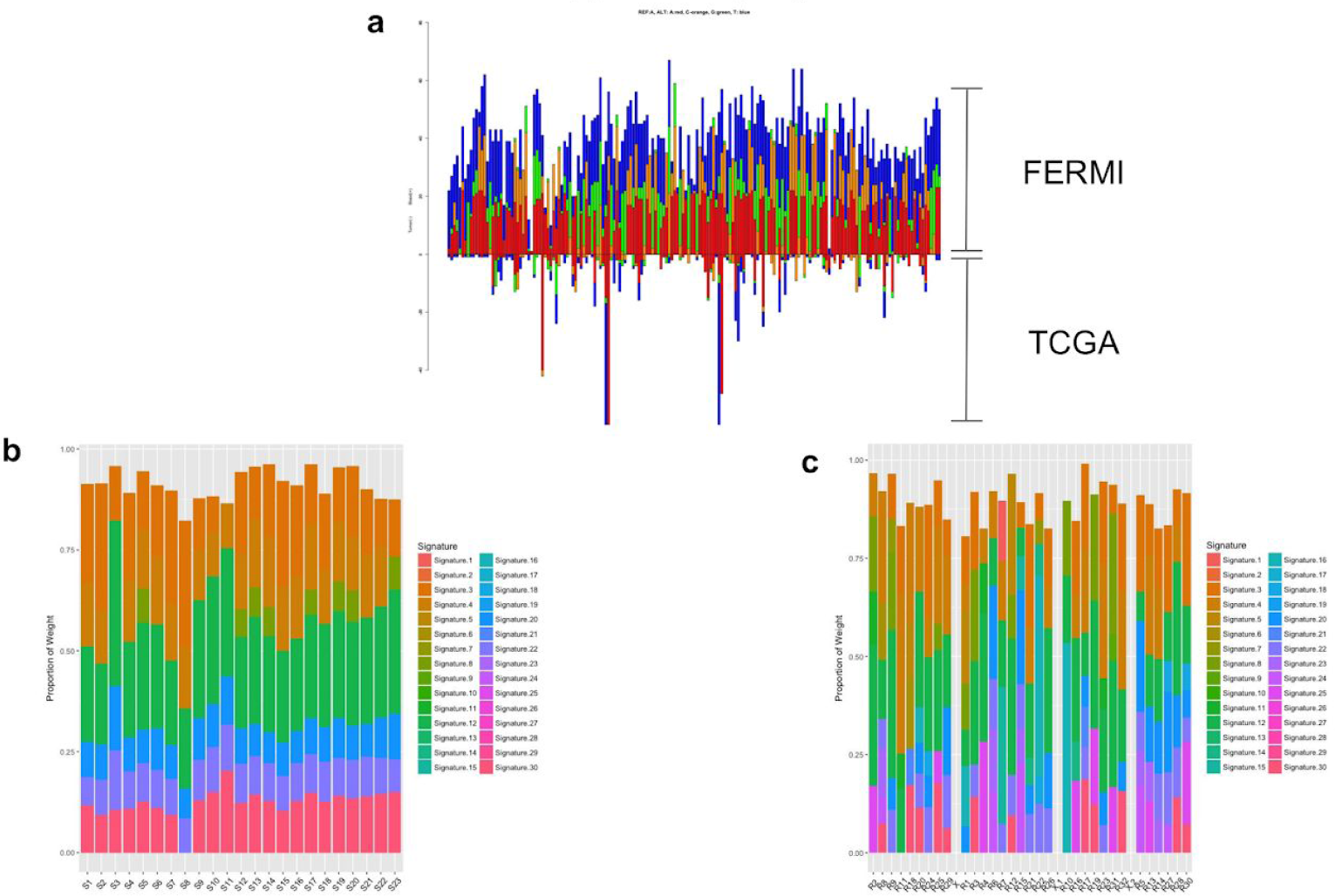
Blood background substitution patterns exhibit previously identified signatures distinct from those in cancers. **a**, We focused on the amplicons in coding regions, and integrated Pan cancer somatic mutation data from exome sequencing in the TCGA to analyze patterns of base substitutions at genomic positions in the target regions which were mutated in tumor genomes and also changed in the background generated for healthy peripheral blood samples. Substitution frequency and substitution patterns were both significantly different between blood and tumors, both at highly mutated sites (mutation count > 10; Chi square test; FDR adjusted p-value <0.05) and across all such sites (Mantel test; p-value < 1e-5), with substitution patterns in tumor genomes being more skewed. It is possible that selection during cancer evolution contribute to the observed patterns. **b**, Integrating trinucleotide contexts of the substitutions, we determined the contributions of different mutation signatures previously identified in the blood gDNA background. Out of 30 previously identified signatures, our data showed overrepresentation of only 7 of them (Signatures 3, 4, 8,12, 20, 22 and 30) across different samples. Out of seven signatures, Signature 12, 3 and 4 had maximum contributions. Signature 3 and 4 are known to be associated with failure of DNA double stranded break repair by homologous repair mechanism and tobacco mutagens respectively, whereas the aetiology of Signature 12 remains unknown. **c**, For the in the blood gDNA background, there was no systematic difference in mutation signatures between amplicons when grouped by their genomic context, and they also showed similar pattern of enrichment of few signatures as compared to others, with signature 12, 3 and 4 having maximum contributions. Signature 12 and 4 exhibits transcriptional strand bias for T>C and C>A substitutions respectively, whereas signature 3 is associated with increased numbers of large Indels.

**Supplemental Figure 4.**
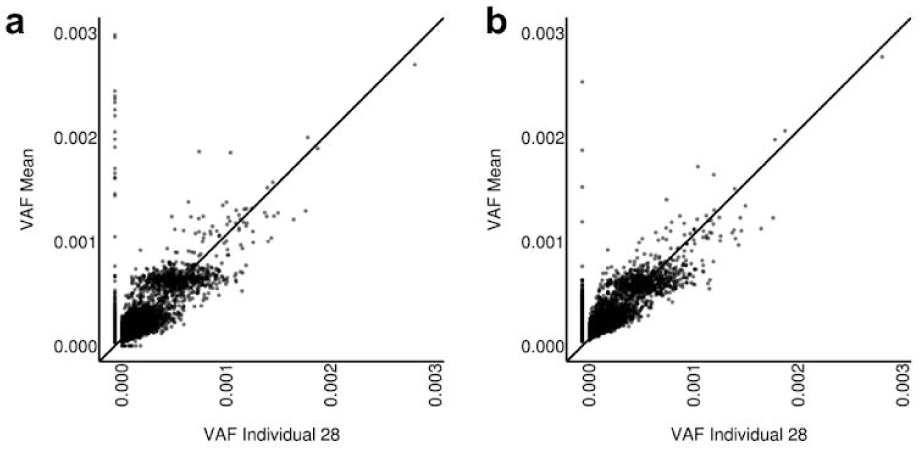
Background patterns are similar across multiple experiments. Background patterns were analyzed across individuals sequenced across different experiments. **a**, Experiment 1 uses a background of 20 healthy individuals from the same experiment. Background VAFs were averaged from these 20 healthy individuals and compared against individual 28 (R-squared = 0.455316, p-value = 0.000000). **b**, Mean background VAFs from 12 individuals sequenced in a different experiment are compared to a resequencing of individual 28 (R-Squared = 0.615327, p-value = 0.000000). Between the two experiments, the background is relatively similar within individual 28, but subtle difference do exist. These subtle differences appear to be somewhat experimentally dependent, and in being so, served as the impetus to only use samples from the same experiment for background derivation where possible.

**Supplemental Table 1.** Trinucleotide context is not sufficient to predict base mutability. To understand how well the trinucleotide context of each unique nucleotide substitution predicts base mutability, all biopsied individuals were split into two groups (Group 1 and Group 2), which were similar in ages. Within these groups, each substitution was sorted by nucleotide and trinucleotide identity. Sorted substitutions were then plotted by their VAF and compared between Group 1 and Group 2. If trinucleotide context is sufficient to predict how often and to what a nucleotide mutates, it would be expected that the comparison between Groups 1 and 2 would result in a uniformly clustered set of variants. If this were the case, the R-squared value would be small as the variant population would not fit a line (for example, the distribution could reflect a round cluster, if substitution frequencies are not correlated in the different sample sets but driven by chance). Alternatively, if factors other than just trinucleotide context were important in determining the mutability of a particular context, it would be expected that variant comparisons between Groups 1 and 2 would strongly adhere to a y=x line and therefore have a high R-squared value. For each context, the R-squared values are shown for the comparisons between Groups 1 and 2. With most comparisons showing a high R-squared value, it is clear that trinucleotide context is not sufficient to predict base mutability.

|  | C>A |  | C>G |  | C>T |  | T>A |  | T>C |  | T>G |  |
| --- | --- | --- | --- | --- | --- | --- | --- | --- | --- | --- | --- | --- |
|  | R-Squared | p-value | R-Squared | p-value | R-Squared | p-value | R-Squared | p-value | R-Squared | p-value | R-Squared | p-value |
| TNT | 0.942188 | 0 | 0.283838 | 0.000008 | 0.942188 | 0 | 0.165416 | 0.001692 | 0.582888 | 0 | 0.582888 | 0 |
| CNT | 0.942336 | 0 | 0.072766 | 0.022908 | 0.942336 | 0 | 0.373684 | 0 | 0.373684 | 0 | 0.838034 | 0 |
| GNT | 0.914237 | 0 | 0.999999 | 0 | 0.914237 | 0 | 0.50136 | 0 | 0.50136 | 0 | 0.595753 | 0 |
| ANT | 0.952932 | 0 | 0.638111 | 0 | 0.952932 | 0 | 0.556291 | 0 | 0.556291 | 0 | 0.488577 | 0 |
| TNC | 0.958693 | 0 | 0.313046 | 0 | 0.958693 | 0 | 0.34038 | 0 | 0.34038 | 0 | 0.622346 | 0 |
| CNC | 0.884787 | 0 | 0.248655 | 0.00042 | 0.884787 | 0 | 0.503585 | 0 | 0.503585 | 0 | 0.690602 | 0 |
| GNC | 0.907913 | 0 | 0.641254 | 0 | 0.907913 | 0 | 0.46375 | 0 | 0.46375 | 0 | 0.999989 | 0 |
| ANC | 0.919144 | 0 | 0.404024 | 0 | 0.919144 | 0 | 0.82306 | 0 | 0.82306 | 0 | 0.871428 | 0 |
| TNG | 0.98003 | 0 | 0.98297 | 0.083316 | 0.98003 | 0 | 0.34038 | 0 | 0.560237 | 0 | 0.658504 | 0 |
| CNG | 0.999966 | 0 | 0.218539 | 0.04357 | 0.999966 | 0 | 0.491406 | 0 | 0.491406 | 0 | 0.77013 | 0 |
| GNG | 0.937916 | 0 | 0.589258 | 0.000012 | 0.937916 | 0 | 0.57064 | 0 | 0.57064 | 0 | 0.77013 | 0 |
| ANG | 0.935708 | 0 | 0.452428 | 0.006004 | 0.935708 | 0 | 0.928586 | 0 | 0.928586 | 0 | 0.627575 | 0 |
| TNA | 0.939775 | 0 | 0.470265 | 0 | 0.939775 | 0 | 0.960272 | 0 | 0.960272 | 0 | 0.350091 | 0 |
| CNA | 0.927451 | 0 | 0.459003 | 0 | 0.927451 | 0 | 0.933907 | 0 | 0.933907 | 0 | 0.999998 | 0 |
| GNA | 0.907147 | 0 | 0.229226 | 0.000016 | 0.907147 | 0 | 0.750916 | 0 | 0.750916 | 0 | 0.739989 | 0 |
| ANA | 0.929315 | 0 | 0.530768 | 0 | 0.929315 | 0 | 0.536645 | 0 | 0.536645 | 0 | 0.408415 | 0 |

